# 3D Molecular Pretraining via Localized Geometric Generation

**DOI:** 10.1101/2024.09.10.612249

**Authors:** Yuancheng Sun, Kai Chen, Kang Liu, Qiwei Ye

## Abstract

Self-supervised learning on 3D molecular structures is gaining importance in data-driven scientific research and applications due to the high costs of annotating biochemical data. However, the strategic selection of semantic units for modeling 3D molecular structures remains underexplored, despite its crucial role in effective pre-training—a concept well-established in language processing and computer vision. We introduce Localized Geometric Generation (LEGO), a novel approach that treats tetrahedrons within 3D molecular structures as fundamental building blocks, leveraging their geometric simplicity and widespread presence across chemical functional patterns. Inspired by masked modeling, LEGO perturbs tetrahedral local structures and learns to reconstruct them in a self-supervised manner. Experimental results demonstrate LEGO consistently enhances molecular representations across biochemistry and quantum property prediction benchmarks. Additionally, the tetrahedral modeling and pretraining generalize from small molecules to larger molecular systems, validating by protein-ligand affinity prediction. Our results highlight the potential of selecting semantic units to build more expressive and interpretable neural networks for scientific AI applications.

## 1. Introduction

Modeling 3D molecular structures is crucial for a wide range of AI applications in drug discovery and material design. Accurate modeling and prediction of molecular properties, such as binding affinity, toxicity, and biological activities, rely heavily on a deep understanding of the 3D conformations and geometric features of the molecules [1, 2, 3, 4].

In recent years, the field of drug discovery has seen a surge of interest in leveraging self-supervised learning techniques to exploit the wealth of unlabeled molecular data available. This is particularly important given the high costs and time-consuming nature of experimental annotation of biochemical properties. By learning from large, unannotated molecular datasets, self-supervised models can capture valuable knowledge embedded in the inherent patterns of the data and acquire meaningful representations that can transfer effectively to a wide range of downstream tasks.

The success of self-supervised learning in other domains, such as natural language processing and computer vision, has demonstrated the critical importance of carefully selecting the appropriate semantic building blocks for model development [5, 6]. By identifying the fundamental units that capture the intrinsic structure and patterns within the data, self-supervised models can learn more expressive and robust representations, which can then be effectively leveraged for downstream tasks.

However, few existing 3D molecular pretraining methods have studied this aspect. Existing 3D molecular pretraining methods fall into two categories: representation-level and structure-level. Representation-level methods aim to enhance 2D molecular representation by leveraging information from 3D molecular structures through contrastive learning [7, 8]. Such methods use 3D molecular structures only at the encoding stage and fail to model inherent structural features through self-supervised training. Structure-level methods address this limitation by developing pre-training tasks of coordinate denoising, where independent noise is added to the coordinates of all atoms in the graph and the model is trained to reconstruct the original atomic positions [9, 10, 11]. However, from a chemical perspective, an atom alone can hardly serve as a functional unit in molecules. Therefore, atom-wise denoising provides limited improvement in the model’s understanding of functional substructures.

In this paper, we focus on this open issue and propose a novel pretraining approach as an initial exploration. Our method, called <u>L</u>ocalized G<u>e</u>ometric <u>G</u>enerati<u>o</u>n (LEGO), treats tetrahedrons within 3D molecular structures as fundamental building blocks, and accordingly designs two pretraining tasks based on these tetrahedral units, thereby facilitating the capture of semantic information and enhancing the learned representations. There are two key conceptual motivations behind this design: Geometrically, the tetrahedron is the simplest polyhedron that can be constructed in 3D Euclidean space, serving as the base case for more complex polyhedra. This structural simplicity and primitiveness aligns with the ubiquity of the tetrahedral motif in chemistry: a central atom along with its one-hop neighbors forms a highly prevalent local structure in molecules, which underlies carbon backbones, functional groups, and more (Fig 1). Therefore, tetrahedrons can be considered an excellent basic semantic unit for 3D molecular modeling from both geometry and chemistry.

**Figure 1:**
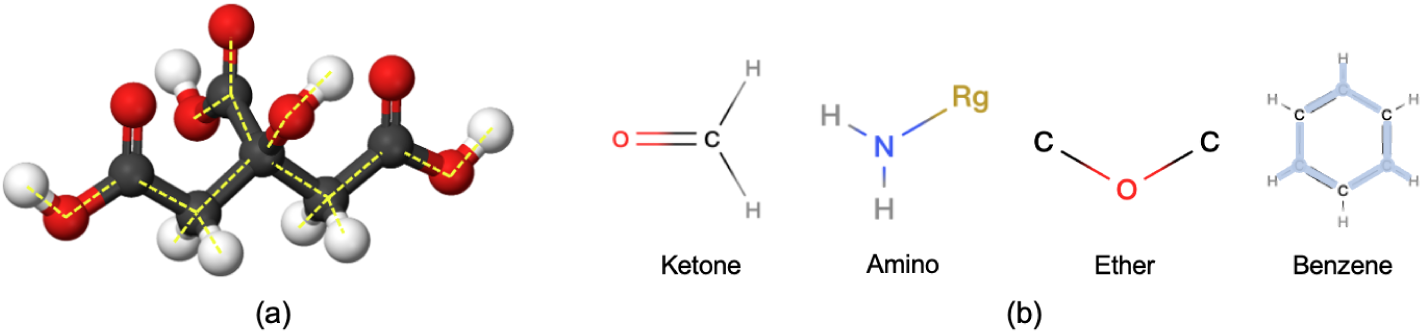
Tetrahedral local structures, comprising a central atom and its one-hop neighbors, represent prevalent structural patterns within molecular data, making them well-suited as fundamental semantic units for self-supervised learning on 3D molecular structures. These patterns include (a) carbon backbones, (b) functional groups, and more.

Inspired by masked language/image modeling techniques [5, 6], LEGO introduces perturbations to tetrahedrons in a 3D molecular structure and learns to reconstruct them during pretraining. Specifically, we begin by segmenting a 3D molecular structure into a non-overlapping assembly of one-hop local tetrahedral structures (LS). We then add noise or apply masking to the local structures segmented. The reconstruction of the perturbed local structures involves two steps: 1) predicting the spherical position of the center atom for each perturbed LS, which provides positional information about local structures and their relationships within the whole molecule; 2) accurately reconstructing the atom arrangements within each perturbed LS, which focuses on learning the intrinsic pattern of the building block itself. By incorporating these global orientation prediction and local structure generation steps, LEGO is able to learn both the overall spatial relationships and the intrinsic geometric patterns of 3D molecular geometry in a self-supervised manner.

**Figure 2:**
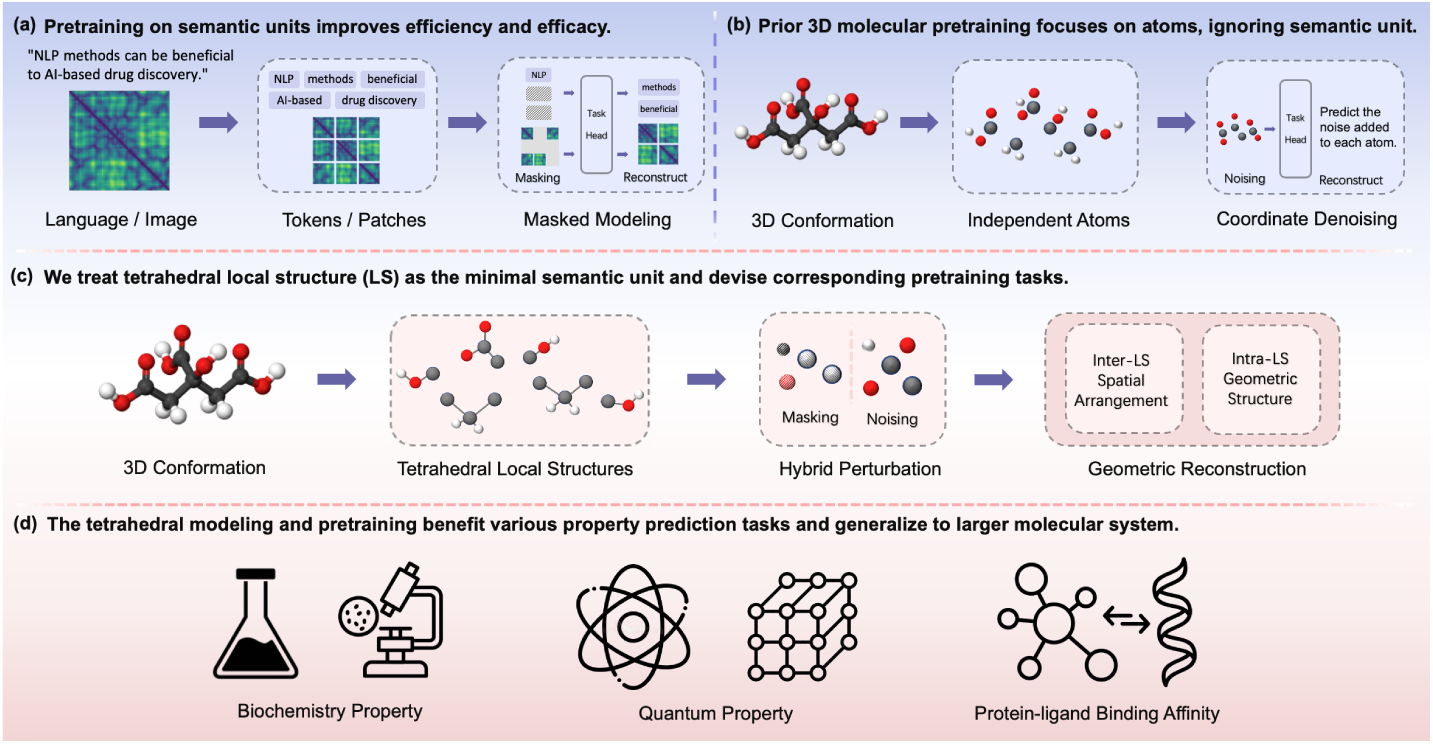
(a) Pretraining on semantic units improves efficiency and efficacy of self-supervised learning, as is widely demonstrated in natural language processing and computer vision. (b) Existing 3D molecular pretraining methods mainly apply atom-level coordinate denoising, which fail to incorporate chemical or geometric semantic prior. (c) In our method, we treat tetrahedral local structures within 3D conformations as the minimal semantic unit, and devise corresponding pretraining task in terms of geometric reconstruction. (d) Demonstrated through extensive experiments, our method enhances a wide range of biochemistry and quantum property predictions, and effectively generalizes to large molecular systems in protein-ligand binding affinity prediction.

We demonstrate the effectiveness of LEGO through comprehensive experiments. Firstly, LEGO significantly enhances representation learning for 3D molecular structures. It sets new state-of-the-art results in 7 out of 8 biochemical property prediction tasks within MoleculeNet [12], and improves performance in 12 out of 16 quantum property predictions within QM9 [13] and MD17 [14]. Secondly, due to the pretraining based on semantic units, LEGO exhibits remarkable generalizability, extending its capabilities from small molecules to larger molecular systems. In the critical task of protein-ligand property prediction, LEGO surpasses baselines by an impressive 17%, demonstrating its substantial potential for practical applications. Finally, we exhibit LEGO’s increased resilience through larger perturbations in pretraining, its enhanced explainability through embedding visualizations, and its broad applicability across experiments on a different backbone structure. These qualities collectively highlight LEGO’s versatility and robustness under diverse scenarios.

## 2. Results

In this section, we detail how LEGO enhances 3D molecular representation learning by conducting downstream fine-tuning on various molecular property prediction tasks. These tasks encompass biochemical and quantum property predictions for small molecules.

Furthermore, we demonstrate LEGO’s remarkable generalizability from small molecules to protein-ligand complexes. Notably, LEGO can predict binding affinities accurately without the structural optimization typically required by conventional methods, showcasing its significant potential for practical applications.

### 2.1. LEGO improves molecule-level property prediction

#### 2.1.1. Biochemistry properties

The biochemical properties of a pharmaceutical molecule, including solubility, permeability, side effects, and toxicity, are closely related to specific semantic substructures such as functional groups. Given that LEGO is designed and pretrained to use tetrahedral local structures as fundamental semantic units in 3D molecular structures, it is expected to demonstrate enhanced capabilities in predicting biochemistry properties. We select biophysics and chemistry tasks in MoleculeNet [12] to conduct the evaluation. We split the datasets based on molecular scaffolds, which is a good approximation of the temporal split commonly used in industry, and adopt an 8:1:1 ratio for training, validation, and testing, respectively.

As shown in Table 1, LEGO achieves state-of-the-art performance on 7 out of 8 benchmark tasks. Notably, LEGO significantly improves predictions of physiological properties (BACE, BBBP), toxicity (Clintox, Tox21) and side effects (Sider), indicating its ability to capture functional semantics within molecular structures. LEGO also performs strongly on FreeSolv and ESOL, where solvation properties correlate with specific functional groups, further demonstrating its effectiveness in capturing structure-function relationships.

**Table 1:**
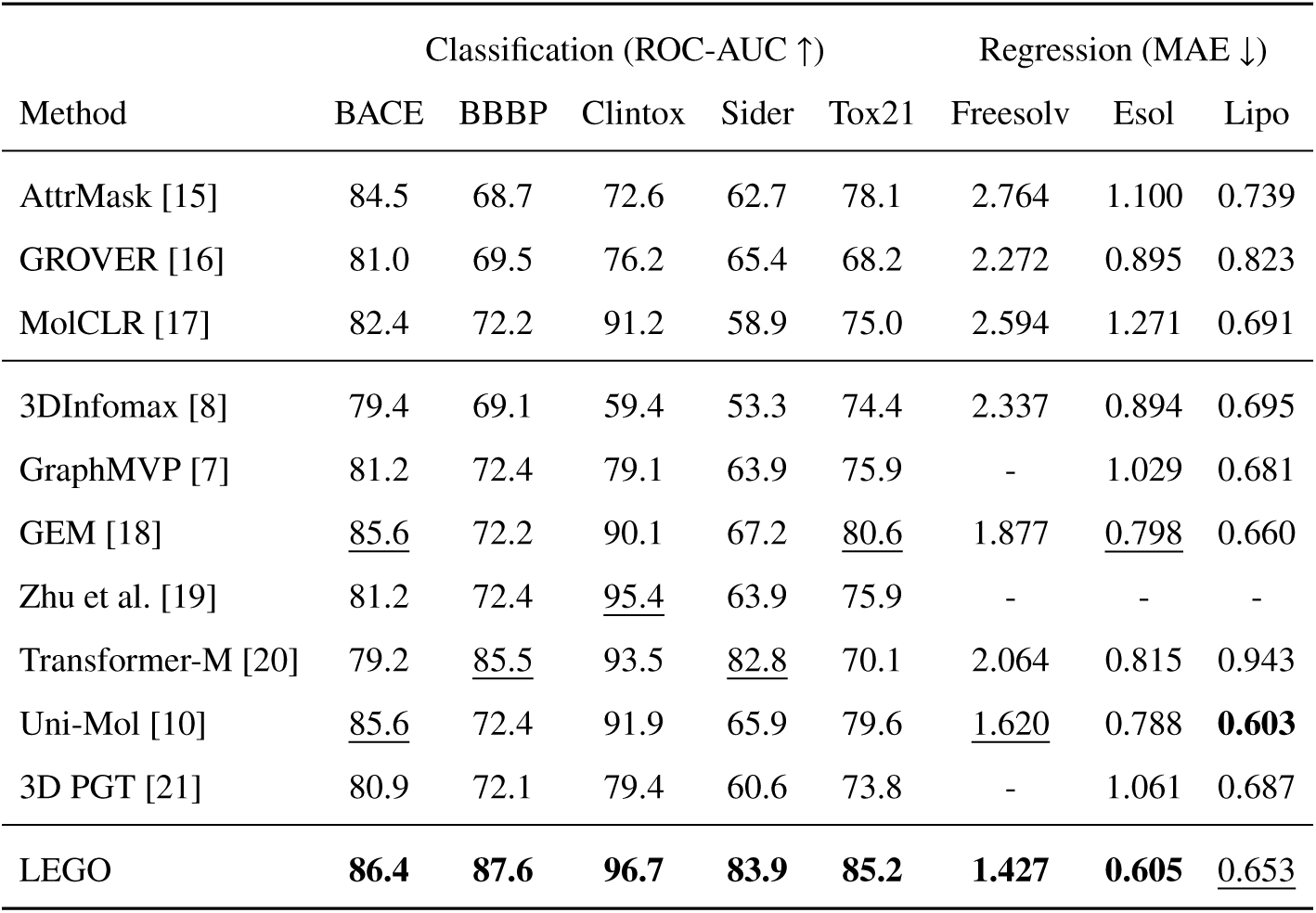
Results on MoleculeNet. We compare our models with existing 2D or 3D molecular pretraining models. The best and second best results are bold and underlined.

Baselines in Table 1 include atom-level coordinate denoising baselines (Transformer-M [20], Uni-Mol [10], and 3D-PGT [21]) and representation-level contrastive learning baselines (3DInfomax [8], GraphMVP [7]). We also include widely-used 2D graph pre-trained models, such as AttrMask [15], GROVER [16], and MolCLR [17], which implement pretraining via masked modeling or contrastive learning. More details regarding the baselines are provided in Section 5.

The superior performance of LEGO over these baselines validates our core hypothesis: proper segmentation of molecular structures is vital for advancing molecular representation learning and functional exploration.

#### 2.1.2. Quantum Properties

Despite focusing on tetrahedral substructures, LEGO effectively captures subtle atomwise interactions within molecular systems, providing a robust framework for accurate quantum property prediction. In this section, we evaluate LEGO’s performance in quantum property prediction using the MD17 [14] and QM9 [13] datasets. The MD17 dataset includes molecular dynamics trajectories for eight organic molecules, emphasizing potential energy and atomic force simulations over time. On the other hand, the QM9 dataset contains about 134k small organic molecules, each composed of CHONF atoms with a single conformer, providing quantum properties such as dipole moment and isotropic polarizability.

Table 2 presents the results for quantum property prediction, including forces from MD17 and eight quantum properties from QM9. Compared to the non-pretrained Transformer-M backbone, LEGO shows performance improvements in the majority of quantum property prediction tasks (12 out of 16). We attribute these improvements to LEGO’s pretext task of localized geometric reconstruction, which enhances the network’s ability to capture subtle atomic interactions and structural nuances critical for accurate quantum property prediction.

**Table 2:** Results (MAE, lower is better) on MD17 force prediction and QM9 property prediction. Compared to the non-pretrained backbone Transformer-M (TM) [20], models pretrained with LEGO exhibit improved performance across most tasks.

| **MD17** | aspirin | benzene | ethanol | MDA | naphthalene | salicylic | toluene | uracil | Avg. ↑ |
| --- | --- | --- | --- | --- | --- | --- | --- | --- | --- |
| TM | 0.096 | 0.243 | 0.085 | 0.0863 | 0.119 | 0.1288 | 0.977 | 0.118 | - |
| LEGO | 0.0935 | 0.241 | 0.079 | 0.0847 | 0.106 | 0.1274 | 0.959 | 0.112 | - |
| $\Delta(\%)$ | 2.67 | 0.83 | 7.59 | 1.89 | 12.26 | 1.10 | 0.88 | 5.36 | 4.2 |

| **QM9** | $\mu$ | $\alpha$ | $\epsilon_{HOMO}$ | $\epsilon_{LUMO}$ | $\Delta\epsilon$ | $R^2$ | ZPVE | $C_v$ | Avg. ↑ |
| --- | --- | --- | --- | --- | --- | --- | --- | --- | --- |
| TM | 0.037 | 0.041 | 17.5 | 16.2 | 27.4 | 0.075 | 1.18 | 0.022 | - |
| LEGO | 0.035 | 0.041 | 16.9 | 16.3 | 27.7 | 0.065 | 1.20 | 0.020 | - |
| $\Delta(\%)$ | 5.4 | 0.0 | 3.4 | -0.6 | -1.1 | 13.3 | -1.7 | 9.0 | 3.5 |

### 2.2. LEGO generalizes to complex-level molecular systems

In molecular-level property prediction, we demonstrate that LEGO effectively captures semantic information in substructures through segmentation and pretraining based on tetrahedral substructures. In this section, we extend this capability from small molecules to larger molecular systems.

We validate this extension using a protein-ligand binding affinity (LBA) prediction task from ATOM3D [24]. The binding affinity prediction measures the strength of the interaction between a small molecule and its target protein. Although the 3D graph of the protein-ligand complex, including the sequences and coordinates of the binding pocket and the ligand, is used as input, this task presents a significant challenge for our pretrained model due to the out-of-distribution nature of such data. To enhance the model’s practicality, the dataset partitions the protein-ligand complexes such that no protein in the test set shares more than 30% (or 60%) sequence identity with any protein in the training set.

Table 3 presents the results of LEGO on ATOM3D-LBA, evaluated using root mean square error (RMSE), which is the primary metric. We also report the Pearson and Spearman correlation coefficients. As shown in the table, LEGO demonstrates a substantial improvement in RMSE over existing baselines, achieving reductions of 17% and 20%. Although LEGO does not achieve the highest correlation coefficients, it ranks second and shows competitive results with the leading value. Notably, LEGO as well as other deep learning methods demonstrate superior performance compared to two conventional methods, CyScore [22] and AutoDock Vina [23], suggesting their potential for practical application. We attribute LEGO’s superior performance to our pretraining approach based on semantic substructures, which enables the model to capture essential interactions within the protein-ligand complex, even when the data distribution differs from the training set.

**Table 3:** Results on the ATOM3D-LBA (Ligand Binding Affinity) dataset. We select structure-based baselines for comparison, and divide them into non-pretraining and pretraining categories. The best and second best results are bold and underlined respectively.

| Method | LBA 30% |  |  | LBA 60% |  |  |
| --- | --- | --- | --- | --- | --- | --- |
|  | RMSE↓ | Pearson↑ | Spearman↑ | RMSE↓ | Pearson↑ | Spearman↑ |
| CyScore [22] | 3.129 | 0.573 | 0.582 | 3.152 | 0.578 | 0.583 |
| Vina [23] | 2.751 | 0.354 | 0.459 | 2.655 | 0.480 | 0.499 |
| Atom3D-CNN [24] | 1.416 | 0.550 | 0.553 | 1.621 | 0.608 | 0.615 |
| Atom3D-ENN [24] | 1.568 | 0.389 | 0.408 | 1.620 | 0.623 | 0.633 |
| Atom3D-GNN [24] | 1.601 | 0.545 | 0.533 | 1.408 | 0.743 | 0.743 |
| Holoprot [25] | 1.464 | 0.509 | 0.500 | 1.365 | 0.749 | 0.742 |
| ProNet [26] | 1.463 | 0.551 | 0.551 | 1.343 | 0.765 | 0.761 |
| SMT-DTA [27] | 1.574 | 0.458 | 0.447 | 1.347 | 0.758 | 0.754 |
| SE(3)-DDM [9] | 1.451 | 0.577 | 0.572 | - | - | - |
| Uni-Mol [10] | 1.434 | 0.565 | 0.540 | 1.357 | 0.753 | 0.750 |
| Frad [11] | <u>1.365</u> | **0.599** | <u>0.577</u> | <u>1.213</u> | **0.804** | **0.801** |
| LEGO | **1.084** | <u>0.596</u> | **0.579** | **1.001** | <u>0.781</u> | <u>0.763</u> |

Other baselines involved in Table 3 include non-pretraining methods like multiple variants of Atom3D [24], Holoprot [25], ProNet [26], and pretraining methods SMT-DTA [27], SE(3)-DDM [9], Uni-Mol [10] and Frad [11].

## 3. Method

In this section, we outline our methodology for 3D molecular structure modeling, which focuses on the use of tetrahedrons as fundamental building blocks. We begin by discussing our rationale for choosing tetrahedrons, followed by detailing the segmentation of molecular structures into tetrahedral units. Finally, we describe our Local Geometric Generation process, which utilizes these units for pretraining. Figure 3 provides an overview of our method.

**Figure 3:**
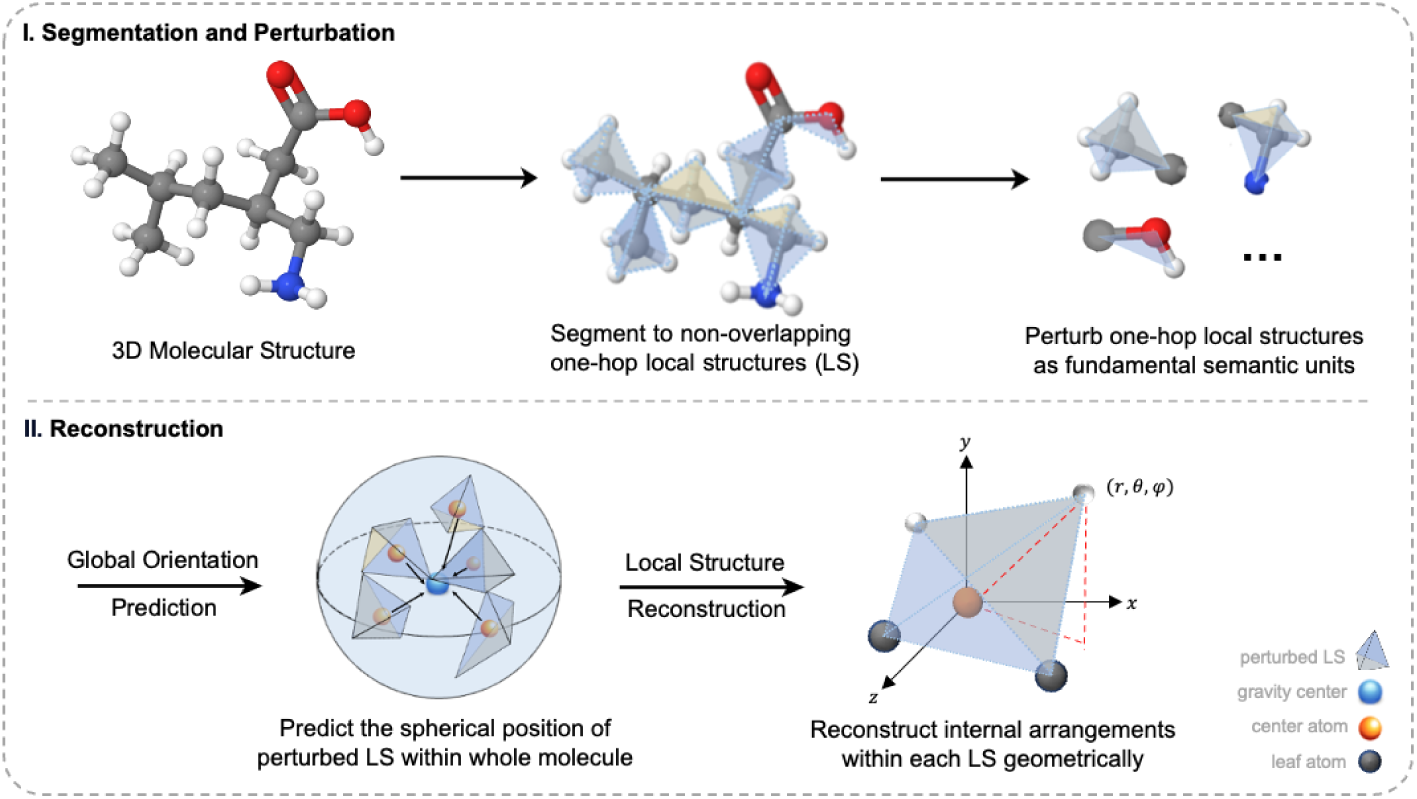
Overview of LEGO. **I.** Based on non-terminal atoms, we segment 3D molecular structures into a non-overlapping assembly of one-hop local tetrahedral structures. We then perturb a portion of the segmented local structures through adding noise and masking features. **II.** We pretrain LEGO by geometrically reconstructing the perturbed local structures in two stages.

### 3.1. Problem Definition

Let *G*_3D_= (*V*, *E*, **P**) denote a 3D molecular structure, where *V* is the atom set {*v*_1_, *v*_2_, *v*_3_, …,}, *E* is the bond set {(*i*, *j*)|*v_i_*and *v _j_* connected}, and **P** ∈ ℝ^|^*^V^*^|×3^ contains 3D Cartesian coordinates for all atoms.

In our approach, we aim to view 3D molecular structures as an assembly of fundamental building blocks, rather than as scattered spatial points. This necessitates a preliminary examination to define these blocks: we need to specify their structural features, identify the critical properties they must possess, and formulate a clear definition for them. Such careful selection is vital for obtaining informative representations through pretraining, as has been widely validated in NLP and CV [5, 6].

### 3.2. Motivation

Semantic units suitable for self-supervised learning must strike a balance between two crucial factors. On one hand, these units need to be structurally insightful, capturing essential details of the local molecular environment and facilitating downstream models for property predictions. On the other, the units should be elegantly simple, avoiding overly intricate or molecule-specific ones that could hinder the method’s generalization across various chemical contexts.

Our proposed solution is to take tetrahedrons – generally one-hop local structures, consisting of a center atom and its one-hop neighbors – as the fundamental building blocks. The geometric simplicity of the tetrahedron, *being the most basic form of polyhedron in 3D space*, makes it an ideal starting point for modeling complex structures. This simplicity is not arbitrary but mirrors the natural occurrence of tetrahedral shapes in chemical compounds, as highlighted in Figure 1. Furthermore, the tetrahedral geometry is fundamental to the architecture of carbon skeletons and various functional groups. Here, the tetrahedral shape, with its valency of four, facilitates the assembly of atoms into complex yet universally applicable molecular structures, *achieving diverse functional complexity.* It allows atoms to connect in a manner that optimizes the use of space, thereby avoiding the physical limitations that might arise from more sophisticated configurations.

In practice, local structures in molecules do not always adhere to a perfect tetrahedral geometry, and our segmentation approach is designed with the flexibility to manage such diversity. In cases where center atoms are connected to fewer than four neighbors, as shown with carbon (C), nitrogen (N), and oxygen (O) in Figure 1(b), we simply treat the ketone, the amino or the ether as a degraded tetrahedra. Additionally, for scenarios where center atoms, such as sulfur (S) and phosphorus (P), form bonds with more than four atoms, our method integrates all neighboring atoms into the local structural analysis. For cyclic compounds like benzene, we select non-adjacent carbon atoms to represent the molecular ring via a synthesis of triangular segments.

### 3.3. Local Structure Segmentation

In this section, we elaborate the segmentation algorithm to divide a 3D molecular structure into tetrahedral local structures as is illustrates in the previous section.

The core idea of local structure segmentation is to ensure none of the segmented results should be overlapped, that is to say, a leaf node in one local structure cannot be the center node in another. We propose to utilize a breadth-first search (BFS) traversal method on the molecular graph to implement the segmentation, which enables the formation of one-hop local connected subgraphs during the traversal process. These subgraphs are then identified as the tetrahedral units. Initially, a node *u*_0_ with a degree greater than two is randomly selected and marked as the center node. Subsequently, its one-hop neighbors are labeled as leaf nodes and enqueued for further BFS traversal. As the BFS queue is processed, a node marked as a leaf node indicates its inclusion in a previously defined local structure and is skipped, whereas an unmarked node (and degree greater than two) is identified as the center of a new local structure. Upon completion of the traversal, two lists of node indices are generated: one representing all center nodes and the other representing all leaf nodes within the segmented molecular structure. While this process does not explicitly record the affiliation relationships between center nodes and leaf nodes, such relationships can be readily inferred from the original graph data. A pseudo code of the segmentation procedure is outlined in Algorithm 1.

**Algorithm 1.**
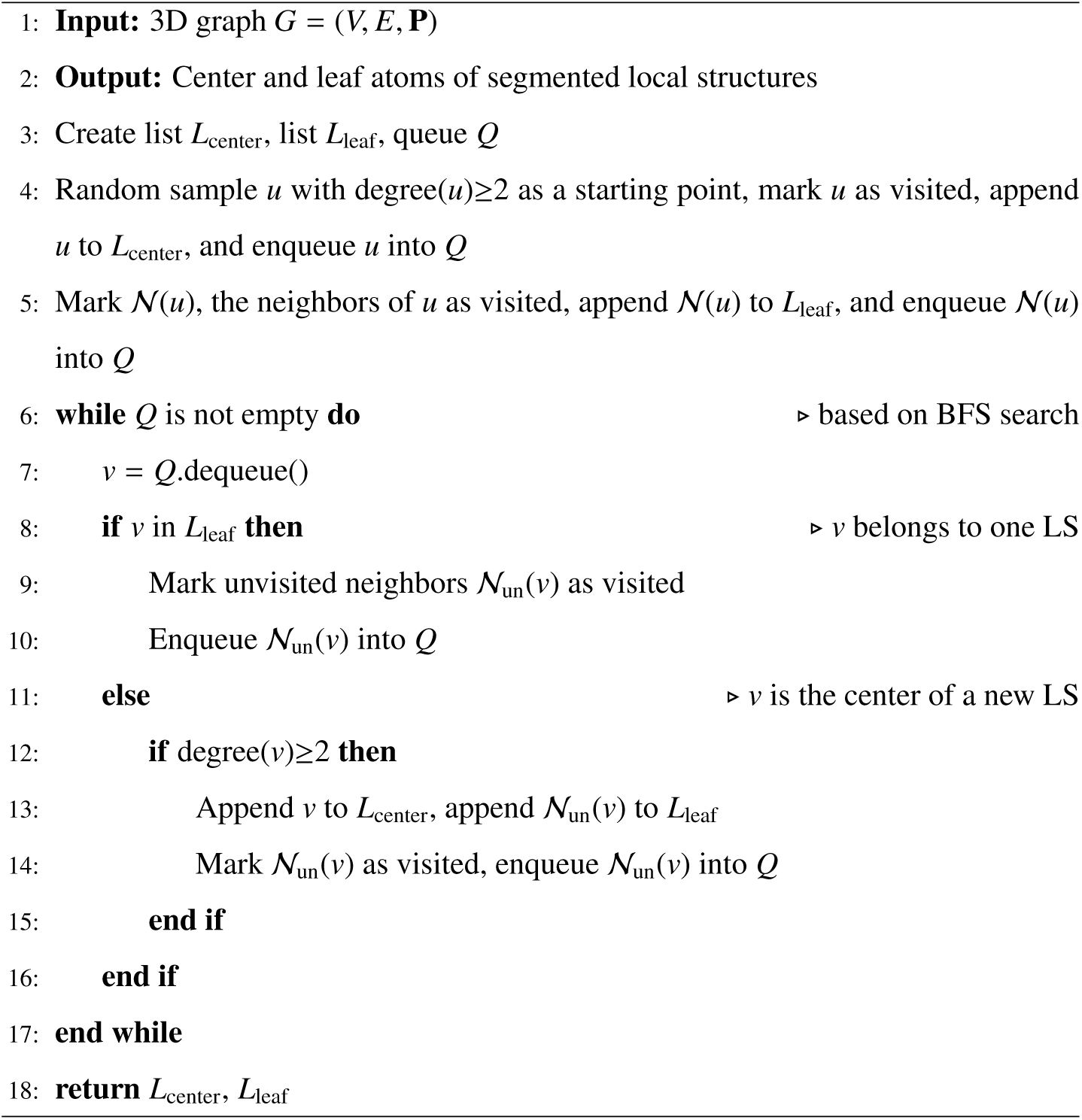
Local Structure Segmentation

**Algorithm 2.**
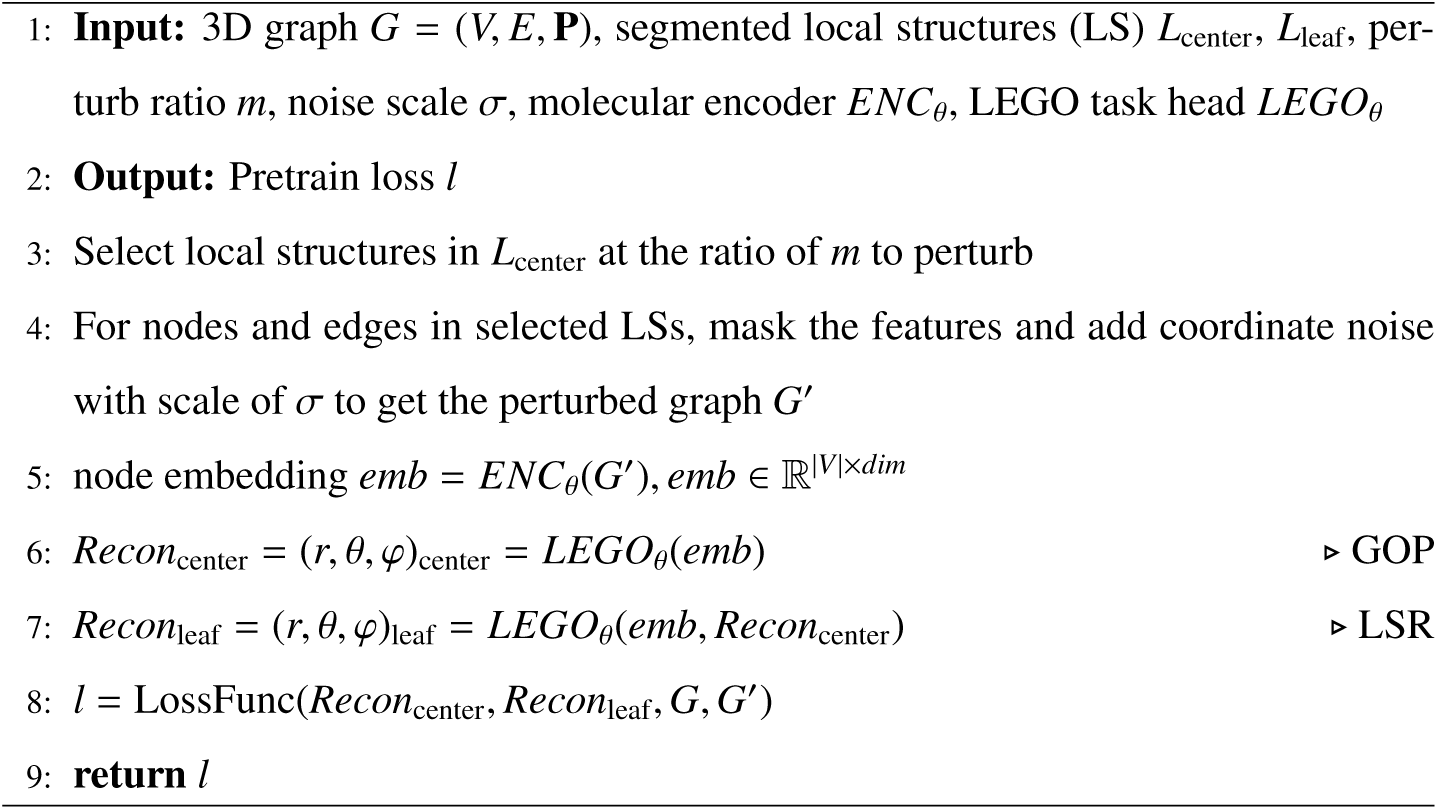
outlines the workflow of the above reconstruction.

For a given graph, our segmentation algorithm can yield different configurations of tetrahedral units due to the randomness in selecting the initial node *u*_0_. This inherent randomness should be viewed not as a limitation, but as a feature to enhance the model’s generalizability and robustness. By varying the central nodes during training, we encourage the model to develop more comprehensive representations, avoiding reliance on a consistent decomposition pattern across pretraining iterations.

### 3.4. Local Geometric Generation

Once the fundamental building blocks have been identified, it becomes imperative to devise a pretraining task that aligns with their specific characteristics. Considering the inherent geometric nature of 3D molecular graphs, a corruption strategy surpassing conventional feature masking is necessary. Furthermore, the design of this pretraining task must ensure the model’s effective capture of structural information during the pretraining phase. Insipired by masked language modeling, we introduce a novel pretraining task termed **L**ocal G**e**ometric **G**enerati**o**n (LEGO), composing of a hybrid perturbation strategy on 3D graphs coupled with a process of geometric reconstruction.

#### 3.4.1. Perturbation on 3D Graphs

Given the segmented results *L*_center_ and *L*_leaf_, we randomly select center atoms at a predefined perturb ratio *m*. Each of these chosen center atoms is associated with a local structure (LS) ready for perturbation, wherein the features of both center and leaf nodes, as well as the features of the edges connecting them, are subjected to masking. In addition, we adopt a noising strategy to the positions of atoms. For the center and leaf nodes within the selected LSs, Gaussian noise is applied to their Cartesian coordinates. By integrating established corruption strategies from both masked modeling and 3D denoising pretraining, we aim to enhance the complexity of the pretraining tasks, prevent information leakage, and foster more effective representation learning directly from 3D structures.

#### 3.4.2. Global Orientation Prediction and Local Structure Reconstruction

To accurately reconstruct perturbed local structures, we focus on two essential elements: the global orientation of each local structure within the entirety of the molecule and the intricate internal configurations of nodes and edges within each local structure. Given a molecular encoder *ENC*_θ_, we can obtain the embedding for each node in the perturbed graph *G*^′^:

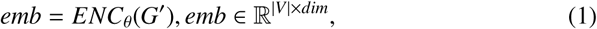

For global orientations, we predict the spherical coordinates of center atoms of the perturbed structures, i.e. radial distances (*r*_center_), azimuthal angles (θ_center_), and polar (φ_center_) angles, originated at the gravity center of a molecule. These geometric elements are essential for defining the precise positioning and orientation of each unit relative to the molecule’s broader architecture.

As for internal geometry reconstruction, we build local frames originated at the predicted center atoms and then predict the spherical coordinates ((*r*, θ, φ)_leaf_) of the leaf atoms within each local frame. In fact, the radial distances *r* represent bond lengths between center and leaf atoms, indicating bond types. While the angular measurements θ and φ encode geometric information and reflect physical and chemical features such as delocalization, steric hindrance, and energy states.

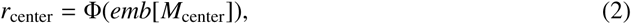

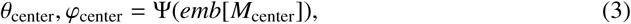

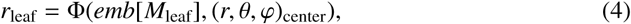

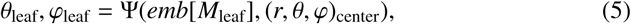

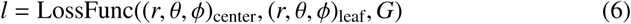

Eqn.(2)-(5) formulates the two-stage reconstruction process, where Φ and Ψ are trainable projection heads for length and angle prediction respectively, and {*M*_center_, *M*_leaf_} ∈ **1**^|*V*|^ are two mask indicators for *V* derived from *L*_center_ and *L*_leaf_, *l* in Eqn.(6) is the final training objective, where LossFunc calculates the loss on lengths and angles between prediction values and ground-truth labels derived from *G*.

## 4. Extended Analysis and Validation

In this section, we perform an extended analysis and validation of our proposed method. This analysis includes ablation studies, an embedding visualization, a probing task, and an evaluation on a different backbone architecture.

The ablation studies help identify the contribution of each component of our method by systematically removing or altering parts of the model. Further, we design a probing task to investigate the model’s internal representations and to assess whether it captures the desired properties and knowledge. The visualization offer an intuitive understanding of how our model makes decisions and validate the motivation of our method. Additionally, we demonstrate the broad applicability of our method by evaluating its performance on a different backbone architecture. Through these combined approaches, we aim to validate our results and provide deeper insights into the effectiveness and functionality of our method.

### 4.1. Ablation Studies

This section evaluates the effectiveness of key components in the LEGO approach, including the structured noise and the hybrid perturbation strategy. We also analyse the influence of hyperparameters during pretraining, i.e. the perturb ratios and noise scales.

#### 4.1.1. Structured Noise vs. Random Noise

Generally speaking, LEGO propose to use structured noise that adds noise only on perturbed tetrahedrons, which distincts it from previous works that add isotropic Gaussian noise on all independent atoms, which we term as random noise [9, 10]. Such structured noise is designed to adhere closely to the inherent geometry of molecular structures, while random noise is less representative of molecular variations among functional groups. As is shown in Figure 4, the random noise can outperfrom non-pretrained model but lag behind LEGO, validating the contribution of the structured noise.

**Figure 4:**
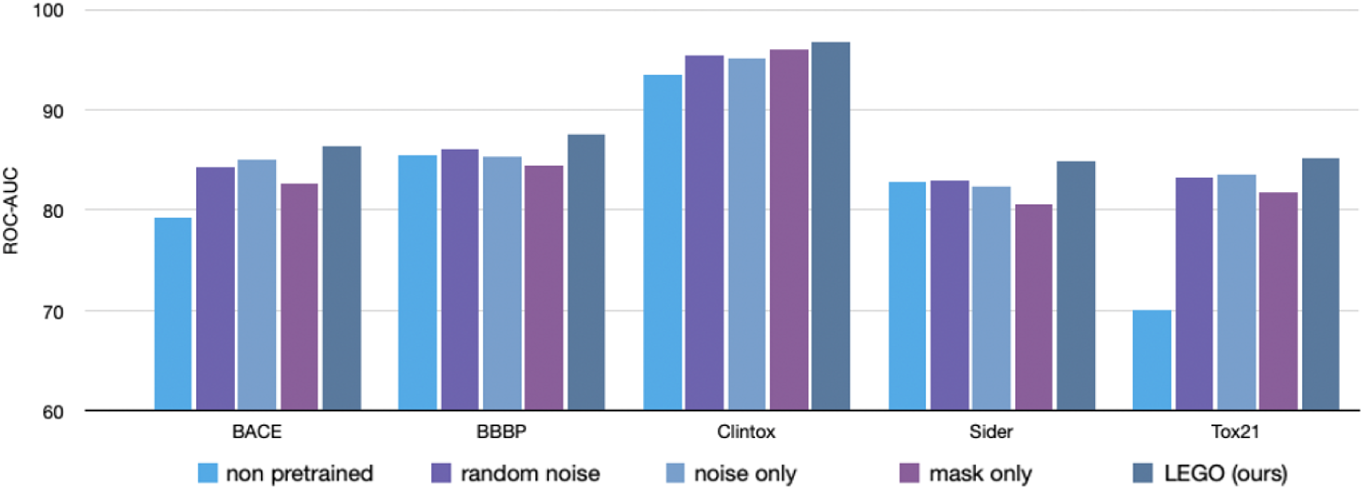
Results of ablation studies on LEGO. The results validate the design of structured noise and hybrid perturbation strategy in the proposed LEGO approach.

#### 4.1.2. Perturbation Strategies on 3D Graphs

In Section 3.4.1, we introduce a hybrid perturbation strategy that integrates both noise addition and feature masking to construct informative pretext tasks for the pre-training process. This section provides a comparative analysis of distinct perturbation strategies on 3D graphs, specifically focusing on noise-only and mask-only strategies.

We hypothesize that the noise-only strategy may lead to information leakage, as the model can infer target predictions directly from the perturbed data. For example, during the local structure reconstruction step, the model might deduce edge lengths from edge type features without considering the context of surrounding coordinates. As for the mask-only strategy, the coordinates remain intact while only the node and edge features are masked. This naive masking task might be less informative or even impair the model due to the significant bias in element type, as is evidenced in previous works [28].

Figure 4 presents the results of these two strategies on MoleculeNet, supporting our hypothesis. The noise-only strategy yields slightly inferior results than its hybrid counterpart. And the mask-only model exhibits the poorest performance among the tested strategies, which is consistent with previous findings. In summary, our hybrid perturbation strategy can effectively increase task complexity, mitigate information leakage and thus enhance the 3D molecular representation learning.

#### 4.1.3. Influence of Perturb Ratios and Noise Scale

**Perturb Ratio.** In our perturbing strategy, we select local structures at a mask ratio of *m* and then perturb and reconstruct these selected structures. When a local structure is chosen, the corresponding center atom, as well as the remaining leaf atoms and edges within that local structure, are all perturbed. This results in actual perturb ratios, denoted as *m^*, that differ slightly from the specified mask ratio *m*. Therefore, it is less intuitive to interpret *m* compared to the mask ratios commonly used for language tokens or image patches. To provide more clarity, we present the corresponding statistics between *m* and *m^* in Table 4.

**Table 4:** Statistics of the perturb ratio *m* on local structures (LS), as well as the corresponding ratios *m^* on all atoms and edges. The figures are means calculated over the entire pretraining dataset, with the values in parentheses indicating the standard deviation.

| $m$ | 0.05 | 0.10 | 0.15 | 0.20 | 0.25 | 0.30 |
| --- | --- | --- | --- | --- | --- | --- |
| LS Perturbed per Molecule | 1.0<sub>(0.2)</sub> | 2.5<sub>(0.7)</sub> | 4.0<sub>(1.0)</sub> | 5.3<sub>(1.3)</sub> | 6.0<sub>(1.4)</sub> | 6.2<sub>(1.4)</sub> |
| Leaf Atoms Perturbed per LS | 3.2<sub>(0.7)</sub> | 3.3<sub>(0.5)</sub> | 3.3<sub>(0.4)</sub> | 3.3<sub>(0.4)</sub> | 3.3<sub>(0.4)</sub> | 3.3<sub>(0.4)</sub> |
| Atoms Perturbed per Molecule | 4.6<sub>(1.5)</sub> | 10.8<sub>(3.8)</sub> | 17.0<sub>(5.4)</sub> | 23.0<sub>(6.6)</sub> | 25.8<sub>(6.8)</sub> | 26.3<sub>(6.7)</sub> |
| Edges Perturbed per Molecule | 3.5<sub>(1.3)</sub> | 8.3<sub>(3.1)</sub> | 13.1<sub>(4.4)</sub> | 17.6<sub>(5.4)</sub> | 19.8<sub>(5.5)</sub> | 20.2<sub>(5.4)</sub> |
| Equivalent Atom Perturb Ratio | 0.16<sub>(0.1)</sub> | 0.36<sub>(0.1)</sub> | 0.57<sub>(0.1)</sub> | 0.77<sub>(0.1)</sub> | 0.87<sub>(0.1)</sub> | 0.90<sub>(0.1)</sub> |
| Equivalent Edge Perturb Ratio | 0.06<sub>(0.02)</sub> | 0.14<sub>(0.03)</sub> | 0.22<sub>(0.04)</sub> | 0.29<sub>(0.04)</sub> | 0.33<sub>(0.04)</sub> | 0.34<sub>(0.05)</sub> |
| Equivalent Token Perturb Ratio | 0.09<sub>(0.03)</sub> | 0.21<sub>(0.04)</sub> | 0.33<sub>(0.05)</sub> | 0.45<sub>(0.05)</sub> | 0.51<sub>(0.07)</sub> | 0.52<sub>(0.08)</sub> |

As we can see from the table, the number of local structures perturbed increases from 1 to 6 as the perturb ratio *m* rises from 0.05 to 0.30 (row 1). A similar increasing trend is observed for the number of perturbed atoms and edges (rows 3 and 4). However, the number of leaf atoms perturbed per local structure (row 2) remains steady at 3.3 with a standard deviation of 0.4, regardless of the changes in the other metrics. This consistent behavior again validates our tetrahedral segmentation approach, confirming that the tetrahedral one-hop local structure is a fundamental and ubiquitous building block within 3D molecular structures.

Figure 5 (left) illustrates the performance of pretrained Transformer-M on MoleculeNet with different perturb ratios (0.05, 0.10, 0.15, 0.20, 0.25, 0.30). The non-pretrained Transformer-M is included as *m* = 0 for comparison. The results show that model performance increases with the perturb ratio initially. However, when the ratio exceeds 0.15, divergent trends emerge across the different tasks. For BACE and Tox21, the model achieves its peak performance at the largest perturb ratio of 0.30. Conversely, the other three tasks exhibit a decline in performance as the perturb ratio is increased beyond 0.15. We attribute this performance divergence to the increasing difficulty of the reconstruction task as the perturb ratio grows. With *m* = 0.15, approximately 60% of all atoms are masked, and this ratio rises to around 80% for *m* = 0.20. Under such heavy perturbations, the model struggles to accurately reconstruct the geometric structures and learn the relevant semantic information during pretraining. Considering these results, we select a perturb ratio *m* = 0.10 for LEGO.

**Figure 5:**
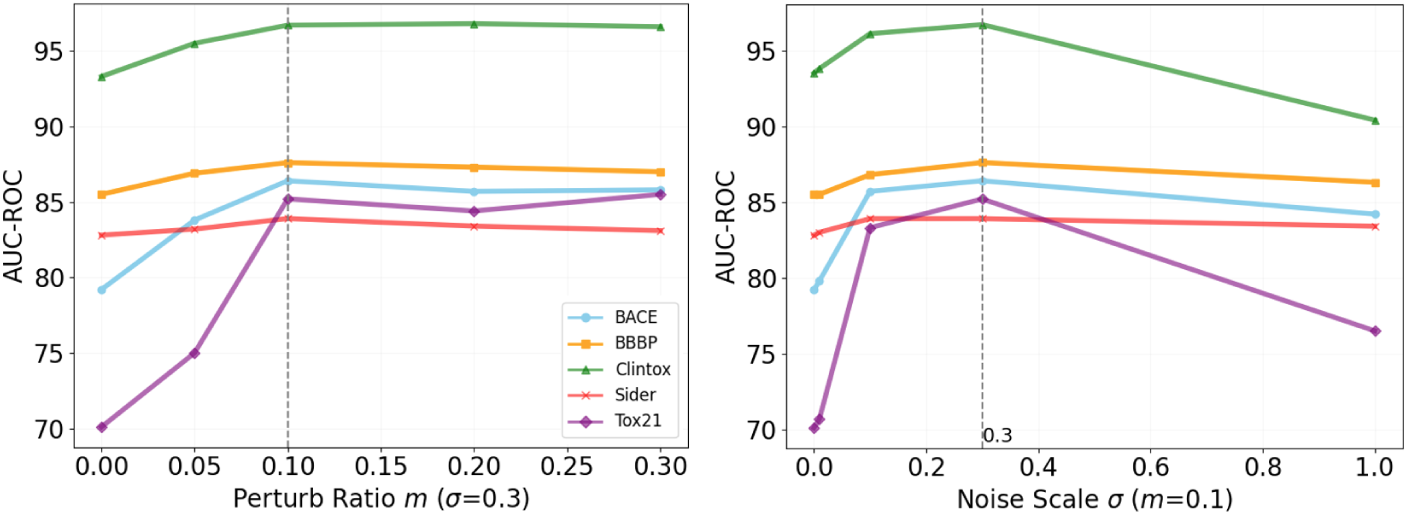
Influence of the perturb ratio and noise scale during pretraining, as measured by the ROC-AUC metric on MoleculeNet classification tasks. The LEGO approach uses a perturb ratio of 0.10 and a noise scale of 0.3, as indicated by the dashed line in respective figures. The right figure shares the same legend as the left one.

**Noise Scale.** The noise scale σ represents the standard deviation of the Gaussian distribution employed to perturb the molecular structures during pretraining. Small perturbation noise can be too trivial for the model to predict, while large noise may break the molecular topology, to the detriment of representation learning.

To select the suitable σ, we evaluated a range of candidate values {0.01, 0.1, 0.3,1.0} Å, as illustrated in Figure 5 (right). The results demonstrate that even the lowest noise scale of σ = 0.01 can bring benefit to the pretrained models. Moreover, models trained with a noise scale of σ = 0.3 were observed to slightly outperform those utilizing σ = 0.1. Notably, this larger noise magnitude exceeds that employed in prior works [11, 29]. We attribute this magnitude advantage to the geometric pretraining tasks, which focus on reconstructing the local tetrahedral structures, rather than relying on naive coordinate denoising. However, when an excessively large noise scale of σ = 1.0 is applied, the pretraining is still found to harm the model’s downstream ability. Based on above experimental results, we select a noise scale σ = 0.3 for LEGO.

### 4.2. Probing Task

To further investigate the model’s internal representations and to assess whether it captures the desired properties and knowledge, we introduce a probing task to evaluate the quality of the atomwise embeddings generated by our model. Specifically, this task aims to assess the ability of the embeddings to accurately identify the functional group to which each atom belongs. Such fine-grained understanding of molecular structures is essential, as it indicates whether the model can effectively leverage the tetrahedral segmentation and local geometric generation pretraining we employed.

We randomly select a subset from the PCQM4Mv2 pretraining dataset, which includes 10k molecules. Each molecule in this subset has its functional groups annotated using the Rdkit toolkit, serving as the basis for our probing task. We omit the more narrowly defined species to simplify the classification process, ultimately formalizing the probing task as a multi-class classification for 50 functional groups.

To conduct this task, we employ a molecular encoder *ENC*_θ_, to generate atomwise embeddings *emb*, based on a uncorrupted graph *G*. These embeddings are subsequently processed by a two-layer MLP designed to classify the atom into one of the functional group categories. This classification step is performed without further training of the encoder θ nor the MLP. The encoder’s parameters, θ, can originate from the model without pretraining, the model pretrained via random denoising, or the model pretrained using our LEGO strategy. The MLP is initialized using the Xavier method to ensure optimal parameter starting values. The prediction results are evaluated using the AUC-ROC metric.

Table 5 displays the results of the probing task. It can be seen that the model utilizing LEGO exhibits enhanced performance in identifying the correct functional groups of atoms, demonstrating advantage over both the unpretrained baseline and the random denoising approach. This result underscores the efficacy of our method in generating embeddings reflecting a deeper understanding of molecular fuctionalities, thereby validating the tetrahedral segmentation and localized geometric generation design.

**Table 5:** Results of probing task on functional group identification.

| Strategy | LEGO | Random | Non pretrained |
| --- | --- | --- | --- |
| AUC-ROC | 0.417 | 0.391 | 0.383 |
| $\Delta(\%)$ | 9.1 | 2.1 | - |

### 4.3. Embedding Visualization

In this section, we apply t-Distributed Stochastic Neighbor Embedding (t-SNE) to visualize the embeddings on two distinct molecular datasets from MoleculeNet. Figure 6 illustrates the t-SNE plots of molecular embeddings generated by LEGO-pretrained and non-pretrained models.

**Figure 6:**
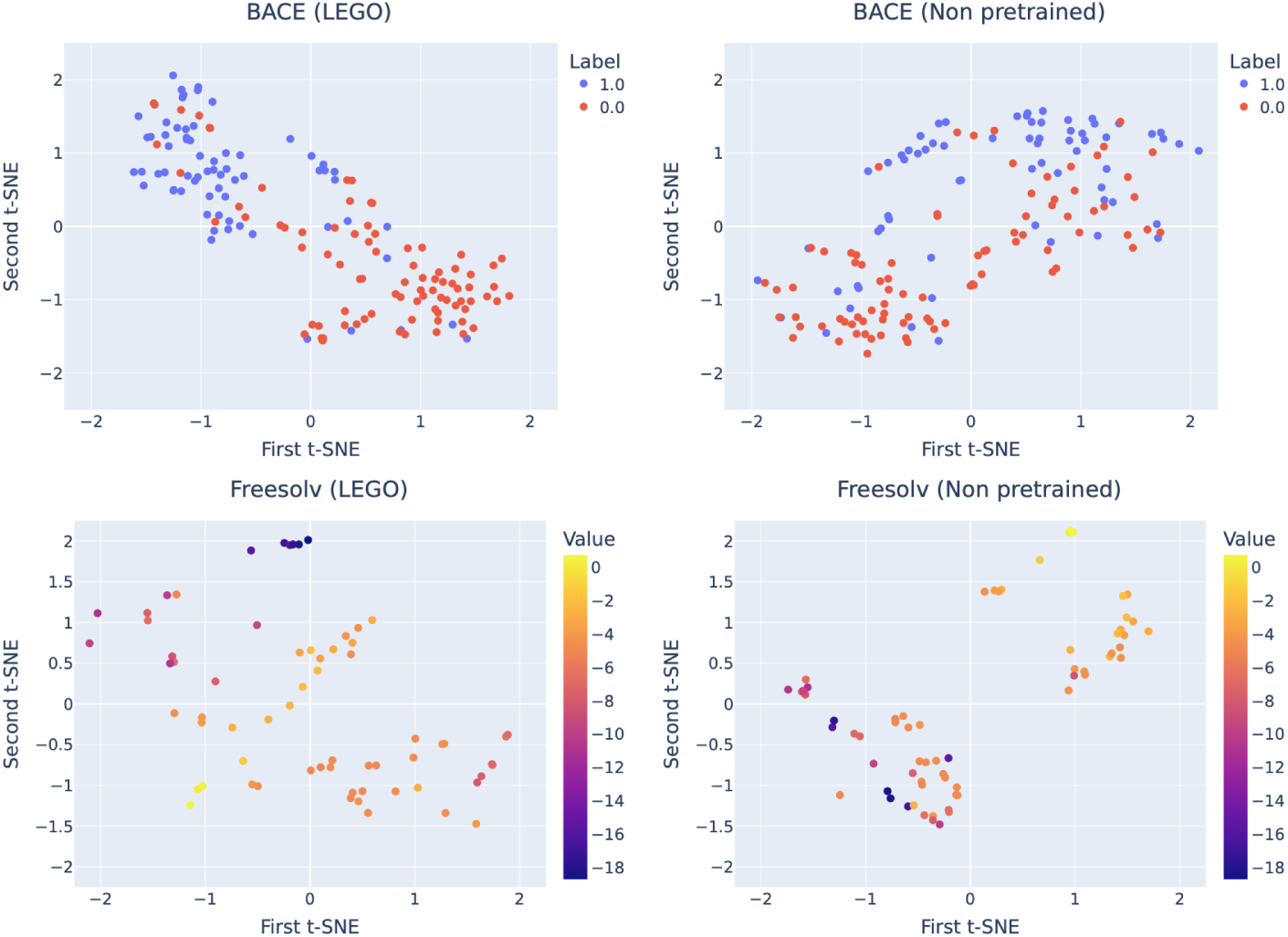
The t-SNE visualization of the embeddings from the LEGO-pretrained and non-pretrained models on two molecular datasets. For the classification task BACE, the pretrained embeddings from distinct classes exhibit enhanced separability, suggesting the deep model has learned more discriminative representations. For the regression task Freesolv, the pretrained embeddings are more uniformly and well-structured distributed across the visualization plane, indicating our method can facilitate the learning of a more organized feature space.

For the BACE classification task, before pretraining, the embeddings of label 0 (represented by red dots) are scattered across the entire plane. In contrast, after pre- training, the embeddings from distinct classes are more effectively separated, suggesting that the LEGO pretraining helps the model learn more informative and distinctive representations. While for the Freesolv regression task, compared to the entangled non-pretrained embeddings, the pretrained embeddings shift smoothly along the continuous labels, suggesting they can lead to accurate predictions in downstream tasks. Further, the pretrained embeddings display a uniformly distributed pattern across the visualization plane, which validates the robustness of the learned representations and demonstrates that the LEGO pretraining helps develop a more coherent and well-structured feature space.

### 4.4. Generalizability of LEGO

In this section, we demonstrate the generalizability of LEGO by applying the proposed pretraining strategy on a different molecular encoder. We choose TokenGT [30] as the new encoder to conduct the experiment. Different from Transformer-M that adds hand-crafted structural encodings onto attention matrix, TokenGT simply treat all nodes and edges as independent Laplacian-augmented tokens and feed them to a standard Transformers. Without graph-specific modifications, TokenGT is still able to achieve promising results in graph learning both in theory and practice.

We first pretrain the TokenGT model using the LEGO approach, and then finetune the pretrained model on the MoleculeNet and PCQM4Mv2 datasets [3]. For the MoleculeNet dataset, we select several classification tasks related to biochemical properties. As for the PCQM4Mv2 dataset, we take the HOMO-LUMO gap as the target. The HOMO-LUMO gap is an important and informative quantum property in biochemistry, and it is present in the PCQM4Mv2 dataset, though it was not utilized as a training objective during the pretraining phase. As is shown in Table 6, LEGO yields performance improvements in both biochemical and quantum property predictions, corroborating the generalizability and versatility of our method.

**Table 6:** Results on TokenGT [30] with and without LEGO pretraining. The results demonstrate the generalizability of LEGO.

| Models | Classification (AUC-ROC) |  |  |  |  | Regression (MAE) |
| --- | --- | --- | --- | --- | --- | --- |
|  | BACE | BBBP | Clintox | Sider | Tox21 | PCQM4Mv2 |
| Non pretrained | 78.3 | 72.2 | 90.2 | 70.9 | 78.5 | 0.0910 |
| Pretrained with LEGO | 81.9 | 74.2 | 94.3 | 72.3 | 83.9 | 0.0817 |
| $\Delta(\%)$ | 4.6 | 2.8 | 4.5 | 2.0 | 6.9 | 10.2 |

## 5. Related Works

The initial focus for 3D molecular modeling is on creating robust neural networks for representations, including Graph Neural Networks (GNN) and Graph Transformers. As the field matured, the emphasis evolved to embrace self-supervised learning approaches to take advantage of large-scale unlabeled datasets.

### 5.1. 3D Molecular Structure Modeling

SchNet [31] first use continuous-filter convolutional layers to integrates pairwise distances between atoms, enabling the capture of local atomic correlations. Following this, DimeNet [32] introduce bond angles and directional elements as geometric features to enhance model performance. SphereNet [33] refine the geometric features by modeling the 3D structures in a spherical coordinate system. In pursuit of embedding 3D equivariance into models, SE(3)-Transformers [34] and NequIP [35] adopt tensor product in message passing and aggregation mechanism.

The exploration extends beyond GNNs, with the transformer architecture for graph-structured data gaining attention. Dwivedi and Bresson [36] pioneers the application of a fully-connected transformer to graphs, employing Laplacian eigenvectors for node positionas. Subsequent works like GRPE [37] and Graphormer [38] introduce structural positional encodings, leveraging node topology, interactions, and 3D distances. Further innovations come from EGT [39] and GraphGPS [40], which propose hybrid models combining message-passing layers with global attention mechanisms. Despite these advancements in directly incorporating 3D features into models, there is an ongoing need for developing pretraining paradigms tailored to 3D molecular structures.

### 5.2. Pretraining on 3D Molecular Structures

Early pre-training methods for 3D molecular structures typically employ distinct encoders for 2D graphs and 3D structures, generating embeddings that undergo contrastive or generative self-supervised learning [7, 8]. These approaches, while valuable, give precedence to 2D representations, thereby ignoring the spatial features and falling short in directly deriving representations from 3D structures.

In contrast, recent advancements have introduced coordinate denoising as a structure-level pretraining strategy, which involves adding specific noise to 3D molecular structures and training neural networks to predict the noise. SE(3)-DDM [9] and UniMol [10] have incorporated Gaussian noise to establish a correlation between coordinate denoising and force field learning. Frad [11] critiques the use of isotropic Gaussian noise for its failure to reflect the anisotropic nature of molecules, proposing instead a fractional noise strategy as a remedy. Zhu et al. [19] adopts a novel strategy within structure-level pretraining by utilizing masked modeling to infer the coordinates of masked atoms from 2D features, moving beyond traditional denoising to bridge modalities. Similarly, GEM [18] and 3D-PGT [21] utilize geometric features, i.e. bond lengths, angles, and dihedral angles, as pretraining tasks, fostering a explicit modeling rather than simple denoising. Building upon these advancements, our work emphasizes a detailed modeling via local semantic units. By doing so, it offers an advanced method for capturing the intrinsic features within 3D data, enhancing both the depth and breadth of molecular structure understanding.

## 6. Discussions and Limitations

Our work represents an initial exploration into the utilization of minimal semantic units for modeling molecular structures. Future research directions could focus on refining the tetrahedral segmentation algorithm for 3D molecular structures, particularly towards functional group-based segmentation. Alternatively, frequency-based algorithms similar to byte pair encoding (BPE) can be applied to this domain.

Compared to the tokenizers commonly used in natural language processing, or the vector-quantized codebooks employed in computer vision, our method does not generate a deterministic codebook for molecular substructures. Further research could explore the establishment of such a deterministic codebook, which could support a wider range of downstream tasks, such as molecular structure generation, as well as enable cross-modal interactions between biological data and other modalities, akin to the thriving developments in language-vision modeling.

## 7. Conclusion

In this paper, we propose a novel approach for self-supervised learning on 3D molecular structures, emphasizing the utilization of fundamental building blocks in molecular pretraining. Our method, characterized by structured denoising techniques, successfully integrates local and global molecular features from the perspective of tetrahedral units, capturing inherent geometry and chemistry features within 3D molecular structures. Through pretraining, our strategy exhibits competitive performance and versatility in predicting diverse molecular properties across biochemical and quantum fields. Further, we demonstrate our method’s generalizability from small molecules to protein-ligand complexes through protein-ligand affinity prediction. These results not only validate the effectiveness of our method but also offer promising directions for advancing the modeling of molecular structures.

## Acknowledgement

This work is supported by the National Science and Technology Major Project (2021ZD0111102).

